# Exogenous sequences in tumors and immune cells (exotic): a tool for estimating the microbe abundances in tumor RNAseq data

**DOI:** 10.1101/2022.08.16.503205

**Authors:** Rebecca Hoyd, Caroline E Wheeler, YunZhou Liu, Malvenderjit Jagjit Singh, Mitchell Muniak, Nicolas Denko, David Carbone, Xiaokui Mo, Daniel Spakowicz

**Affiliations:** Division of Medical Oncology, The Ohio State University Comprehensive Cancer Center; The Pelotonia Institute for Immuno-Oncology, The Ohio State University Comprehensive Cancer Center - James Cancer Hospital, and Solove Research Institute; Department of Biomedical Informatics, The Ohio State University College of Medicine

## Abstract

The microbiome affects cancer, from carcinogenesis to response to treatments. New evidence suggests that microbes are also present in many tumors, though the scope of how they affect tumor biology and clinical outcomes is unclear. A broad survey of tumor microbiome samples across several independent datasets is needed to identify robust correlations for follow-up testing. We created a tool to carefully identify the tumor microbiome within RNAseq datasets and then applied it to samples collected through the Oncology Research Information Exchange Network (ORIEN) and The Cancer Genome Atlas (TCGA). We showed how the processing removes contaminants and batch effects to yield microbe abundances consistent with non-high-throughput sequencing-based approaches. We sought to establish clinical relevance by correlating the microbe abundances with various clinical and tumor measurements, such as age and tumor hypoxia. This process leveraged the two datasets and raised up only the concordant (significant and in the same direction) associations. We identify associations with survival and clinical variables that are highly cancer-specific and relatively few associations with immune composition. Finally, we explore potential mechanisms by which microbes and tumors may interact using a network approach. *Alistipes*, a common gut commensal, showed the highest network degree centrality and was associated with genes related to metabolism and inflammation. The exotic tool can support the discovery of microbes in tumors in a way that leverages the many existing and growing RNAseq datasets.

**Statement of Significance:** The intrinsic tumor microbiome holds great potential for its ability to predict various aspects of cancer biology and as a target for rational manipulation. Here, we describe a tool to quantify microbes from within tumor RNAseq and apply it to two independent datasets. We show new associations with clinical variables that justify biomarker uses and more experimentation into the mechanisms by which tumor microbiomes affect cancer outcomes.

## Introduction

The microbiome affects many aspects of human health, including cancer. Interactions with the immune system affect immune cell activity and the levels of systemic inflammation, and thereby various disease states (1). For example, gut microbiome samples collected before treatment with immune checkpoint inhibitors could predict tumor response months later (2–4). Intrinsic tumor microbes have been established for years in a relatively small number of cancers. However, recent studies identified their presence in a broader set of histologies and the ability to use high-throughput sequencing data for their study. Understanding the directionality of causal relationships may yield new approaches for cancer treatment

Causal connections between tumor microbes and cancer outcomes remain relatively few. For example, *Bacteroides fragilis* biofilms on colon polyps have been found to secrete a toxin that directly damages DNA and leads to colon cancer (5). In other cases, *Helicobacter pylori* secretes molecules that elicit an inflammatory cascade that has been shown to drive tumorigenesis in gastric adenocarcinoma (6). The fungal genus *Malassezia* was shown to drive pancreatic ductal adenocarcinoma growth by activating the C3 complement pathway (7). Broader surveys include a recent bacterial ribosome amplification-based approach applied to six types of tumors, which found many microbes (8). Hundreds of negative controls and paraffin-only blocks were sequenced to establish the background signal and system contamination. Further, fluorescence in situ hybridization and immunohistochemistry validated the presence of microbes in many locations in tumor tissues and cells. This seminal work opened the door to studying how the microbes’ provenance and effects on cancer outcomes, such as by leveraging high throughput sequencing datasets.

A recent survey of cancer tissue whole-genome shotgun sequencing data from the publicly available data resource TCGA that used a similar approach to the exotic pipeline (described here) found bacteria and viruses in all tumors tested (9). Further, these exogenous sequences were cancer-specific, and exogenous sequences in blood could predict the type of cancer. Provocatively, this was even true in early-stage colon cancer samples, where no circulating tumor sequences were detectable in the blood samples. This powerful observation was lacking by (i) focusing on DNA-based observations, which may be more prone to contamination, (ii) excluding fungi, which have been shown to affect several cancers, (iii) not validating on an independent dataset, and (iv) not testing other correlations that might confound the association with tumor type, such as the microenvironment.

We extend this previous work with a computational pipeline we call exotic for exogenous sequences in tumors and immune cells. The process involves stringent alignment parameters and a series of filtering steps. We show how validation on two large, independent datasets finds many microbes with concordant behavior in several tumors.

## Results

### Description of the dataset

We processed tumor RNAseq from 480 samples through the exotic pipeline (**Table 1**). These included thoracic (n=202), sarcoma (n=118), gastro-intestinal (n=104), renal (n=20), other genito-urinary (n=20) and cutaneous (n=16) cancers. The characteristics of the patients from which these samples were derived varied by BMI (one-way ANOVA, p-value < 0.001), age (oneway ANOVA, p-value < 0.001), and gender (one-way ANOVA, p-value = 0.045). Patients did not vary by the fraction alive at the last follow-up. We retrieved the most similar cancer types from TCGA, including lung adenocarcinoma and squamous cell carcinoma (LUAD, LUSC), colon adenocarcinoma (COAD), rectal adenocarcinoma (READ), sarcoma (SARC), kidney renal cell carcinoma (KIRC), bladder urothelial carcinoma (BLCA), and skin cutaneous melanoma (SKCM). The TCGA samples showed similar demographic characteristics, except for the fraction metastatic, which was higher in the ORIEN data (p-value < 0.001).

**Table 1.** Data summary. The number of samples and demographic information for tumor RNAseq generated by the Oncology Research Information Exchange Network. Each cancer type showed differences with respect to body mass index, age at the time of collection, gender, and the sample preservation method.

|  | CUT | GI | GU | RCC | SAR | THO | p |
| --- | --- | --- | --- | --- | --- | --- | --- |
| n | 16 | 104 | 20 | 20 | 118 | 202 |  |
| BMI at Collection (mean (SD)) | 34.43 (8.72) | 29.06 (6.18) | 30.54 (8.98) | 34.37 (7.68) | 29.08 (6.03) | 27.03 (5.82) | <0.001 |
| Age at Collection (mean (SD)) | 59.34 (10.78) | 59.14 (11.72) | 60.86 (12.22 ) | 56.27 (11.68) | 56.06 (15.99) | 63.39 (10.04) | <0.001 |
| Gender = Male (%) | 12 (75.0) | 62 (59.6) | 16 (80.0) | 15 (75.0) | 62 (52.5) | 108 (53.5) | 0.045 |
| Vital Status = Deceased (%) | 7 (43.8) | 46 (44.2) | 10 (50.0) | 7 (35.0) | 52 (44.1) | 90 (44.6) | 0.965 |
| FFPE = Frozen (%) | 14 (87.5) | 98 (94.2) | 13 (65.0) | 17 (85.0) | 102 (86.4) | 132 (65.3) | <0.001 |

### Description of the exotic pipeline

We designed the exotic pipeline to broadly but conservatively identify microbes present in the tumors and remove the technical artifacts and contaminants from the dataset (**Figure 1A**). Exotic first maps raw reads with quality scores (FASTQs) to the human reference genome, with a second alignment pass following the standard workflow of TCGA and other large-scale sequencing efforts. Next, exotic aligns the unmapped reads to a wide range of non-human genomes, including bacteria, archaea, viruses, fungi, and a subset of other eukaryotes. Next, exotic filters contaminants in two phases: statistical filtering and literature matching. Exotic removes taxa whose species-level relative abundances significantly correlate with input RNA concentration (10). Further, exotic removes a list of organisms commonly found in sequenced negative controls (11), sparing taxa with strong literature precedence for host interactions (9). Finally, the outputs are normalized to remove technical artifacts. The majority of the analyses rely on intersecting results across multiple datasets in a “discovery” and “validation”-type approach. Here we rely on data from the Oncology Research Information Exchange Network (ORIEN), which has the benefit of robust clinical data to support the analyses (**Table 1**).

**Figure 1.**
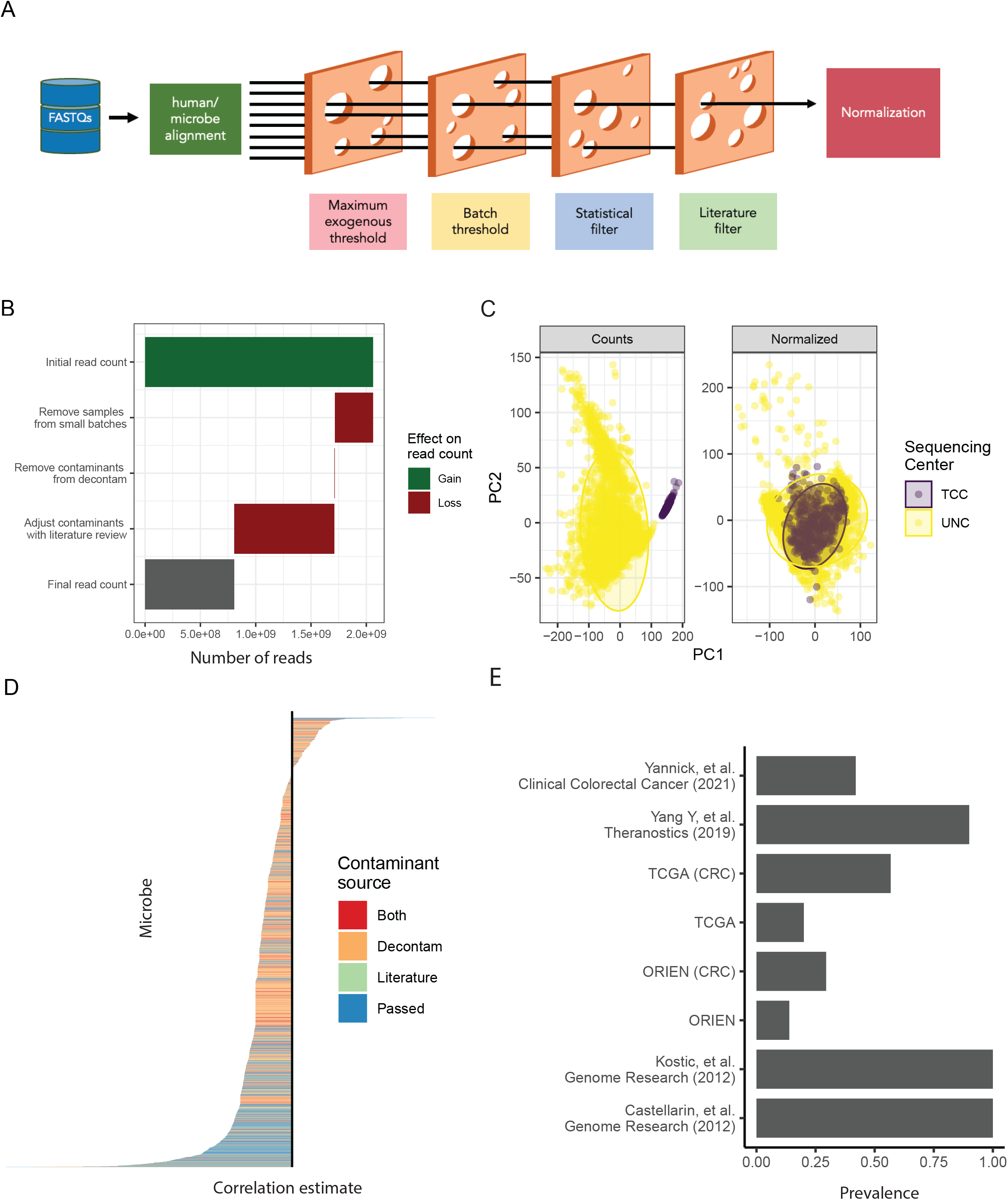
Summary and validation of the exotic pipeline. **(A)** Schematic of the exotic pipeline, showing the process of aligning raw RNAseq fastqs to databases for human and microbial identification, followed by filtering to remove contaminants. Filtering steps include the removal of samples with a high percentage of exogenous reads and samples from small batches that prevent contaminant checks, as well as reads that align to microbes found to be contaminants by statistical or literature review-based filtering. The remaining samples’ microbe counts are normalized by VOOM-SNM. **(B)** The loss of reads at each step of the exotic pipeline. **(C)** Normalization removes batch effects from the sequencing center from decontaminated counts. **(D)** Microbe-level filtering removes taxa that align with input RNA concentration and are consistently found in negative controls. Microbes with strong literature precedence as commensals are returned. **(E)** Comparison of the prevalence of Fusobacterium found by exotic compared to literature values captured by non-RNAseq-based approaches, with a much higher prevalence appearing in colorectal samples compared to all cancer types.

Exotic discards roughly half of the non-human reads in processing (**Figure 1B**). A relatively small fraction of the reads is lost in the statistical filtering step, though these reads represent a large fraction of the total microbes (**Figures 1B, C**). By contrast, the literature-based filtering removes a large fraction of the reads but relatively few taxa (**Figures 1B, C**). After processing and normalization, technical artifacts related to the sequencing center and sample fixation method (formalin fixation, fresh frozen) are removed (**Figure 1C**).

### Association of the tumor microbiome with survival and clinical features

Having identified, filtered, and normalized the microbial abundances, we tested relevance to human health by exploring associations with clinical features and outcomes. First, we target the association of microbes with overall survival across the entire dataset and in each cancer individually. Stratifying samples by the presence of each microbe at every taxonomic level revealed hundreds of significant associations across the ORIEN and TCGA datasets (**Figure 2A**). Following a discovery and validation type approach, we intersect the results to identify the most concordant associations by significance and whether there was a positive or negative effect on survival. Few microbes are associated with overall survival, and very few across more than one cancer type, suggesting cancer specificity. In cases where a microbe was associated with the entire dataset, it was most often associated with one other cancer, which likely drove the effect in the context of the larger dataset. For example, the *Streptomyces* strain CdTB01 is associated with lower overall survival in KIRC, LUAD, and lung tumors and across the entire dataset in aggregate (**Figure 2B, Supplemental Table 1**). This *Streptomyces* strain was later found to have a high betweenness centrality in a network constructed from microbe gene correlations (**Supplemental Table 3**). *Acinetobacter calcoaceticus* presence is associated with reduced overall survival across all tumors in aggregate, with a similar hazard ratio between the ORIEN and TCGA datasets.

**Figure 2.**
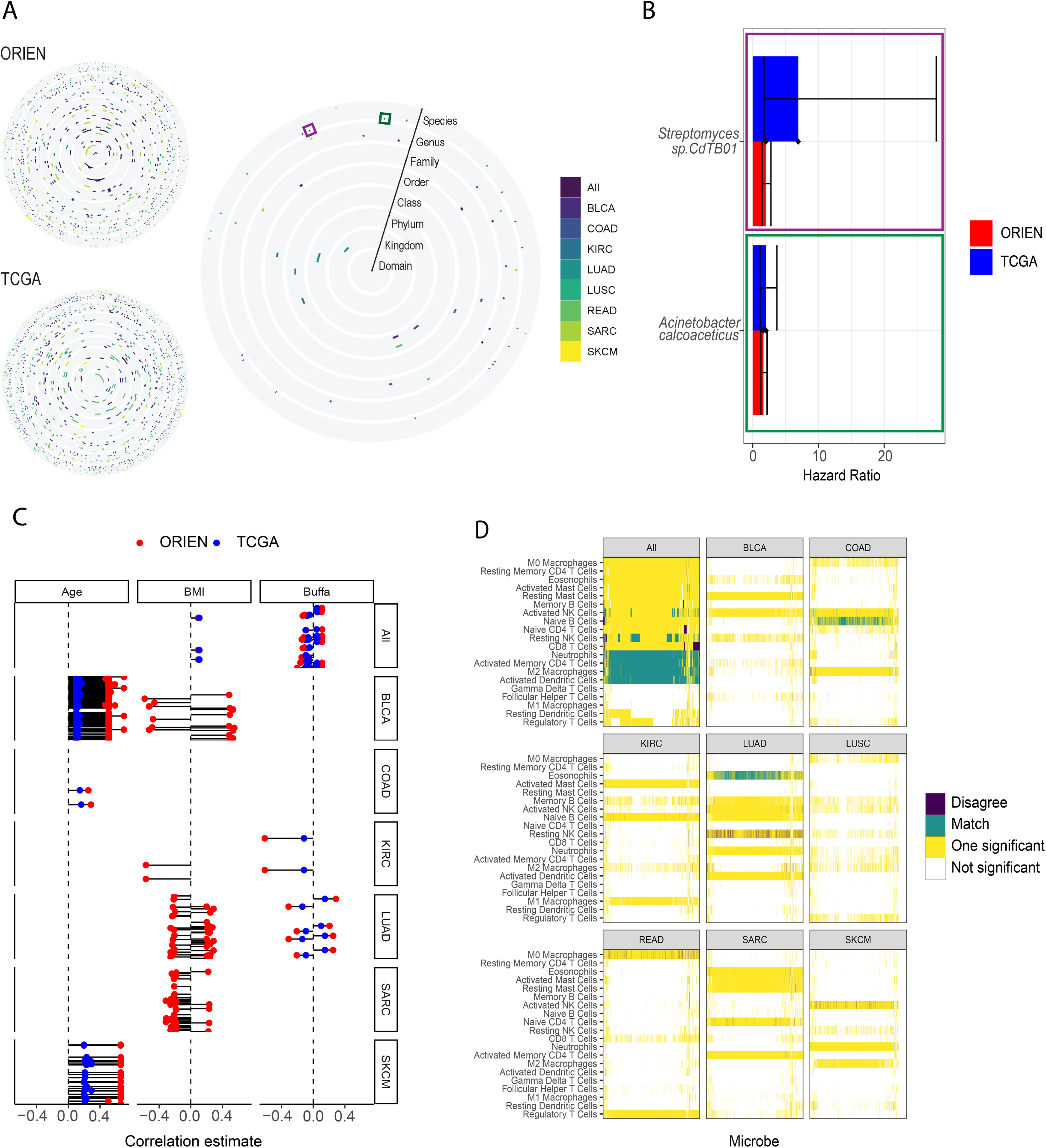
Estimating clinical relevance of observed tumor microbes through agreement between two independent datasets. **(A)** Survival analyses by cancer stratified by microbe presence/absence at every taxonomic level. Colors indicate a significant association with survival. The ORIEN and TCGA analyses separately show many significant associations, whereas few are consistent (significant and affecting survival in the same direction) across both datasets. **(B)** Two example taxa show consistent hazard ratios between the two datasets. **(C)** Microbes correlate with representative clinical variables and an expression signature of the tumor microenvironment (Buffa hypoxia score). Shown are consistent correlations (significant and in the same direction) for ORIEN and TCGA data, except for BMI, which is missing in TCGA for many cancers. **(D)** Correlation of microbes with deconvolved immune cell abundances within each cancer.

The relative abundance of microbes correlated with some, but not all, patient and tumor characteristics. Moreover, we observed enrichment of correlations in certain cancers instead of evenly distributed across the dataset (**Figure 2C**). Again, this analysis relied on observing in both datasets that the correlation was (a) statistically significant and (b) in the same direction. Age correlated with many microbes in breast (BLCA) and skin (SKMC) tumors but not in sarcomas (SARC), lung adenocarcinomas (LUAD), or renal cell carcinoma (KIRC). Few microbes correlated with age in colon adenocarcinoma (COAD). The significant correlations were positive, with an increased relative abundance of a microbe associated with increased age. By contrast, BMI correlations were positive and negative (**Figure 2C**). However, similar cancer specificity was observed, with significant correlations restricted to BLCA, LUAD, and SARC cancer types. Only in the case of BMI do we show significant correlations for ORIEN-only, as the TCGA dataset is missing many values. Finally, we show an association with an expression signature of hypoxia (12). Again, we observed cancer specificity, with bi-directional correlations in LUAD and the entire dataset and only negative correlations in KIRC (**Figure 2C**). Finally, we observed associations between microbe and immune cell abundances estimated by deconvolution. No immune cells significantly correlated with microbes in both datasets and directionally agreed across all cancer types. Neutrophils, activated CD4 memory T-cells, M2-type macrophages, and activated dendritic cells significantly correlated and agreed across the entire dataset. However, only naive B cells and eosinophils significantly correlated and agreed in COAD and LUAD, respectively. Microbes are significantly associated with immune cells in opposite directions in the case of resting NK cells in LUAD and M0 macrophages in READ. More correlations agreed between the two datasets than disagreed, particularly when surveyed across the entire dataset.

### Tumor microbiome-host interactions through gene-microbe networks

To explore the mechanisms by which microbes associate with clinical outcomes such as survival and hypoxia (**Figure 2**), we used a network-based approach to relate microbe abundances with gene expression. We correlated all microbes and all genes across all samples separately in each dataset. The correlation direction agreed in nearly 75% of the cases, with a small fraction significant in both (**Figure 3A**). We then constructed a network drawing edges by the most extreme significant and agreeing correlations (**Figure 3B**). We ranked microbes and genes according to their degree-centrality (**Supplemental Table 2**). The microbe with the highest degree-centrality, *Alistipes*, had edges to nearly 900 genes (**Figure 3C**). *Alistipes* also had the second highest betweenness centrality of all microbes (**Supplemental Table 3**). Other microbes with high degree-centrality included species of *Bifidobacterium, Escherichia, Salmonella*, and *Fusobacterium*. The genes connected to *Alistipes* represented pathways related to adipogenesis, xenobiotic metabolism, interferon-gamma response, and others (**Figure 3D**). The genes connected to Bifidobacterium shared many pathways with genes connected to *Alistipes* but included hypoxia and UV response (**Figure 3E**).

**Figure 3.**
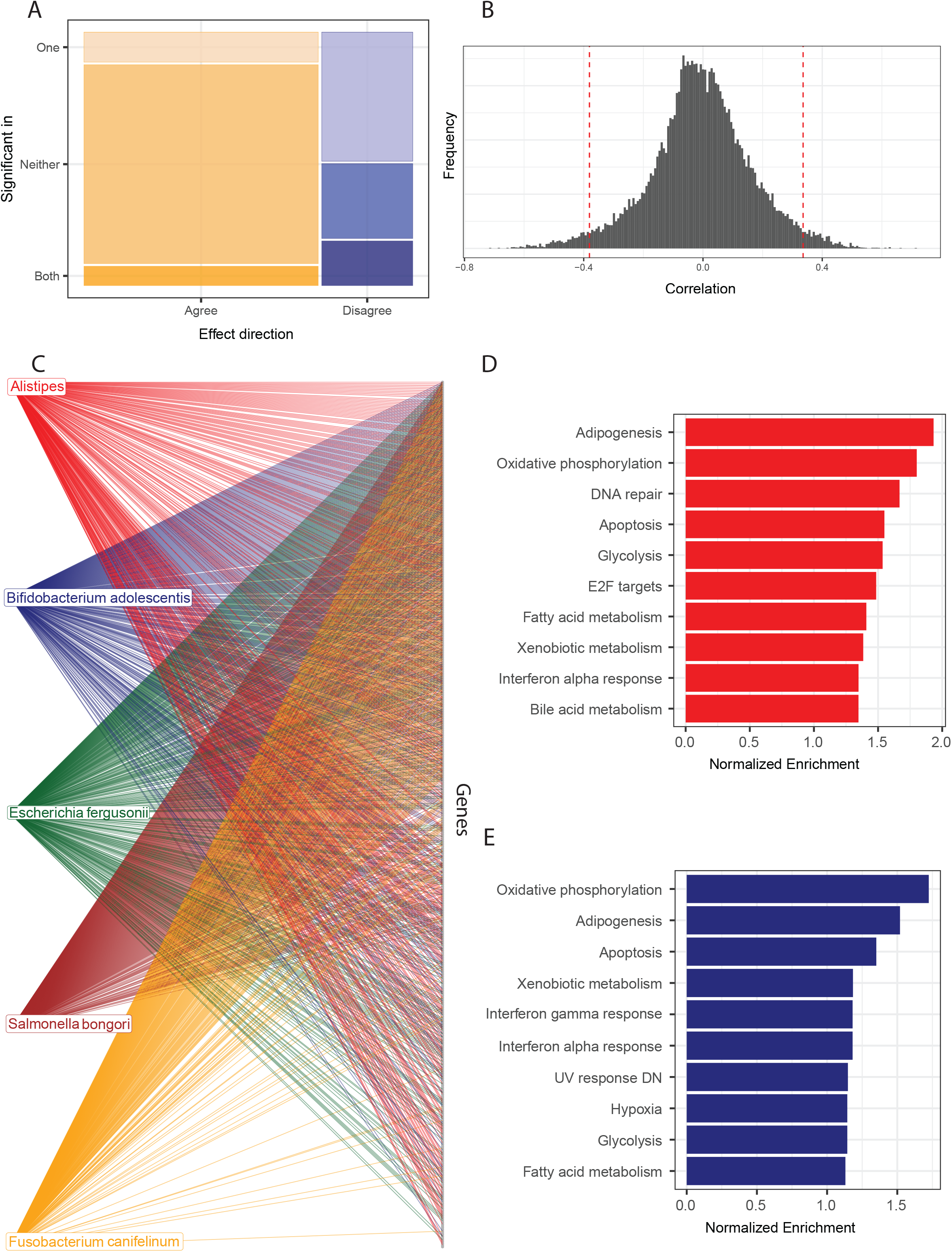
Exploration of mechanism through microbe-gene interaction networks. **(A)** Summary of the correlations between all microbes and all genes, selecting only the correlations significant in both datasets and where the effect direction agrees. **(B)** Consistent correlations are further filtered for the 5% most extreme values. **(C)** Consistent and extreme correlations are then used to build a network. The top taxa by degree centrality are shown, which include those with precedence for affecting cancer outcomes (*Bidobacterium*), as well as microbes that have only been described in the gut (*Alistipes*). **(D)** *Alistipes*-associated genes are enriched in inflammation-related pathways (interferon alpha response) and xenobiotic metabolism. **(E)** *Bifidobacterium adolescentis*-associated genes are enriched inflammation, metabolism, and hypoxia pathways.

## Discussion

The exotic pipeline identifies low abundance microbes within tumor biopsies using well-established alignment tools followed by a series of filtering steps. These include sample-based and microbe-based filtering, which utilizes fractions of microbial sequences in the tumors, the correlation with technical variables, and then lists of common contaminants and common commensals to arrive at a dataset with more than a 50% reduction in observed microbes. Despite these efforts, we show large differences between analogous datasets from TCGA and ORIEN that may be due to contamination or other differences between the datasets, e.g., the type of treatment patients received. Therefore, the analyses presented here rely on concordant results for both TCGA and ORIEN datasets with respect to the direction of effects and their significance. With this framework, we explored the role of microbes in clinical outcomes and their association with clinical variables and immune cells. Further, we built a pan-cancer microbe-gene interaction network to survey mechanisms by which microbes interact with tumors.

The methods used here are consistent with others that support the presence of microbes in many tumors. For example, Nejman *et al*. performed extensive amplicon and immunohistochemistry-based assessments of tumor samples, including nearly a thousand controls. The result was detectable bacteria in over half of breast and bone cancers and fewer in melanoma and other cancers (8). By contrast, computational-only methods that used one dataset reported microbes in all tested tumors (9). Others used multiple datasets in the contamination identification step, showing that microbial sequences in peripheral blood mononuclear cell samples from the same patients showed contaminant-like properties (13). As many datasets lack paired blood samples, we instead showed how the positive-associated based intersection method is an analogous method to reduce the false discovery rate.

Notable differences with our strategy include focusing on RNA rather than DNA-based sequencing. This selection was a deliberate attempt to reduce contamination, as DNA may persist longer in the environment than RNA purification and sequencing apparatus (11,14). In addition, we expanded the database of potential microbes to include fungi, owing to recent reports of fungi increasing tumor growth in pancreatic ductal adenocarcinoma (7) and affecting response to radiation in models of breast cancer and melanoma (15). Despite these differences, we found relatively well-characterized microbes in prevalences consistent with previous reports. Namely, *Fusobacterium* was found in roughly 30% and 60% of colorectal tumors for ORIEN and TCGA, respectively, but only 10% of non-colorectal tumors. This enrichment in colorectal tumors is consistent with other studies that used non-RNAseq-based methods, such as amplicon or microscopy. The consistency with previous reports gave us the confidence to explore associations with clinical data.

The presence or abundance of tumor microbes is associated with a variety of clinical variables, including overall survival, age, and BMI, and tumor features such as the level of hypoxia and immune cell composition. While many microbes and taxonomic levels were associated with survival in the ORIEN and TCGA datasets separately, a much smaller number was observed in both. Moreover, no microbes are consistently associated with survival in more than one cancer. This suggests microbe-cancer specificity that may provide a biomarker for cancer type or treatment outcomes. These observations are consistent with previous reports that showed machine learning models could predict cancer types with high accuracy (9). This association with survival does not take into account the treatment patients receive. The two datasets were collected over different periods (2006-2013 for TCGA and 2003-2018 for ORIEN) when the dominant treatment began to shift to immunotherapy. Differences have been shown to affect survival in the context of chemotherapy (16), radiation (15), and immunotherapy (2–4,17,18). Therefore, even tumor microbes consistently identified between the two datasets could show different results with respect to overall survival.

Rank-based correlations with age and BMI also showed cancer-specificity, but in a different form than for survival. For each cancer, either there were many associations found consistently between the two datasets or few. Age correlated with microbes across both datasets in BLCA, COAD, and SKCM cancers but not in KIRC, LUAD, and SARC or across all cancers. Gut barrier function is maintained with healthy aging (19), which is consistent with significant correlations across some, as opposed to all, cancers. Further investigation is needed to identify the cause of the association with this subset of cancers.

Similarly, BMI is associated with BLCA, KIRC, LUAD, and SARC cancers, but not the others. A high BMI has been identified as a risk factor for several cancers, including COAD. In this analysis, unlike any others described here, we depart from the typical format of only showing agreement between the two datasets. BMI, as well as many other clinical variables that have demonstrated importance in cancer, are largely missing from the TCGA clinical data. We, therefore, chose to call attention to a behavior similar to the age-based correlations using the ORIEN dataset alone and to emphasize the need for robust clinical information. Similar specificities were observed with the hypoxia and immune cell abundances, where microbes are associated with a fraction of cancers and of immune cells.

We sought to explore the mechanisms by which microbes and tumors interact using a networkbased approach. By filtering to the strongest correlations, using them to build a network, and then using network features to identify the nodes with the most edges, we ranked microbes by their network importance. Several microbes with established interactions with tumors were among the nodes with a high degree centrality. For example, members of the genus *Bifidobacterium* correlated with genes in pathways related to interferon signaling and hypoxia. Bifidobacterium is a common gut commensal that was shown to affect the response to immune checkpoint blockade via the secretion of inosine, which is taken up by immune cells via the adenosine receptor A2AAR (20). In the tumor microenvironment, high levels of A2AAR ligands have been associated with hypoxia (21). The most connected microbes was *Alistipes*, which is a known gut commensal but without demonstrated effects in tumors. *Alistipes* connected to genes involved in many fundamental processes in the cell, including metabolism. Further research is needed to verify if *Alistipes* has these effects in the tumor.

This work is limited in several ways. First, while we rely on consistency in two independent datasets, there is no orthogonal experimental validation of the presence of microbes or their associations. Second, some analyses, such as the association with overall survival, rely on assumptions about the presence of an organism being important rather than some measure of quantity or location. Third, not all analyses could be confirmed across datasets because of missing clinical values. This includes treatment information, and there are many examples of microbes affecting outcomes conditional on some treatment regimen (e.g., □-Proteobacteria affecting response to treatment with gemcitabine). Finally, many tumor samples are collected years before the patient dies, and the turnover rate for most intrinsic microbes in tumors has not been established, so the microbes observed here may not be present for long.

Exotic is a pipeline for identifying microbes in the context of human-dominated high-throughput sequencing datasets, such as tumor RNAseq. By processing and intersecting analyses in two large, independent datasets, we further solidified the consistent presence of microbes in tumor samples. In addition, we observed new correlations with age, BMI, and features of the tumor microenvironment such as hypoxia and immune cell abundance. The microbes may interact with hundreds of genes, affecting pathways such as metabolism and hypoxia. This is another example of the value of large, publicly-available data repositories such as TCGA and how they can yield new information many years after they are generated. In the case of exotic, several new lines of evidence allow the re-analysis of these data, including (i) more complete microbial genome databases, (ii) thorough contaminant profiling datasets, and (iii) smaller-scale validation studies that increase the confidence in low abundance organisms. Additional work is needed to validate these observations and identify the mechanisms by which they may affect cancer outcomes.

## Methods

### Description of the dataset and data generation methods

Data processing, contaminant removal, and normalization. Bulk RNAseq samples with corresponding expression data were obtained from ORIEN and TCGA. Samples from TCGA were identified and downloaded using the R package “GenomicDataCommons” and restricted to samples from patients’ primary tumors. TCGA projects included LUAD, LUSC, COAD, READ, SARC, THCA, SKCM, KIRC, and BLCA. These samples were then classified using Kraken2 with Bracken and a custom database built using the available libraries archaea, bacteria, fungi, human, plasmid, protozoa, UniVec, and viral. From here, the R package “decontam” was used to identify contaminants within ORIEN and TCGA samples (10). These contaminant lists were revised using the literature-reviewed list from Poore *et al*. before contaminants were removed from the count tables. The counts were then normalized using VOOM-SNM, treating cancer type as a biological variable to be preserved and the sequencing center and sample preservation method as technical variables whose variance should be removed. The relative abundances of all human and microbe portions of the samples are then calculated. All prevalences are calculated from rarefied, decontaminated counts.

### Correlation/other modeling methods, including Survival methods

All microbe relative abundance correlations with age, Buffa hypoxia score, BMI, and all immune cell fractions were tested using Spearman’s ranked correlation test. Unnormalized expression counts were deconvolved to immune cell fractions using CIBERSORT(22). Overall survival was tested using microbe prevalences that were not otherwise normalized in Kaplan-Meier curves. These tests were performed independently in the TCGA and ORIEN datasets. The results were then compared to find microbes that were concordant in both datasets with respect to significance and the direction of effect between the groups.

### Network construction and analyses

All microbes were correlated with all genes using Spearman’s ranked correlation test in both the TCGA and ORIEN datasets. The ORIEN results were filtered to keep only correlations that had the same direction of effect as the corresponding correlation in the TCGA data. The top and bottom 2.5% correlations were then used to define the edges of a network as the most extreme correlations. The degree centrality of this network was calculated as the number of connections formed with each node, and the betweenness centrality was calculated using the R package igraph (23). Microbes with a high degree centrality and literature significance were chosen for pathway analysis with R package FGSEA (24) using the hallmark pathway database.

### Comparison across analyses

Microbes were given indicators for each of the correlation analyses in which they were found to be consistently significant in the TCGA and ORIEN datasets. These indicators were then summarized in a table with the degree and betweenness centrality information from the network included with it.

### Reproducibility and data access

All data are available through NCBI GEO through the bioproject PRJNA856973. The tool, and scripts to run all analyses presented here, are available in: https://github.com/spakowiczlab/exotic.

